# Response of photosynthetic efficiency to short-term fluctuating light and cold in tomato

**DOI:** 10.1101/2024.07.02.601750

**Authors:** Laavanya Rayaprolu, Keshav Jayasankar, Mark G. M. Aarts, Jeremy Harbinson

**Affiliations:** Laboratory of Genetics, Wageningen University and Research, Droevendaalsesteeg 1, 6708PB Wageningen, The Netherlands; Laboratory of Biophysics, Wageningen University & Research, Stippeneng 4, 6708 WE, Wageningen, The Netherlands

**Keywords:** tomato, fluctuating light, temperature changes, photosynthesis efficiency

## Abstract

Climate-resilient crops are crucial for meeting global food demand and increasing crop productivity. Photosynthesis, a crucial process, is impacted by environmental changes such as temperature and irradiance. Photosynthesis and stomatal opening often lag behind these changes, resulting in a loss in Light Use Efficiency (LUE). Temperature variations also affect photosynthesis, with a decrease below the optimal threshold resulting in a decrease in photosynthetic efficiency. To enhance photosynthetic LUE, understanding plant responses to environmental changes is essential. This study examines the short-term responses of four tomato genotypes to irradiance fluctuations using chlorophyll fluorescence and the effects of transient cold stress. The results show genotype-to-genotype variation in the maximum quantum efficiency of PSII, the kinetics of the quantum efficiency of PSII’s response to step changes in irradiance, and steady-state values of ΦPSII, which is used as a stand-in for photosynthetic efficiency. The control conditions were measured at 24°C and the cold stress conditions at 14°C. The fact that ΦPSII responds dynamically to step decrease and increase in irradiance and how cold impacts these responses illustrates the way tomato genotypes are impacted by cold stress. It also reveals how the genotypes adapt to cold exposure and recover once the cold stress is reversed.

**Highlight:** This study investigates the adaptation and recovery of four tomato genotypes to irradiance fluctuations and transient cold stress, highlighting the importance of climate-resilient crops for food demand.

## Introduction

Starting with an estimated 7.6 billion people in 2021, overall global demand for all agricultural production is expected to rise by 1.1% year until 2050. The annual consumption of crops, the use of those crops for the production of livestock and fish, and all losses (waste and spoilage during the production, storage, transportation, and manufacturing of food and crops) are also impacted by the economic disparities across countries. As a result, a challenge in the coming decades will be to assure long-term food security for the world’s growing and increasingly affluent population by boosting crop production through sustainable agriculture practices while also maintaining soil health and environmental quality (Noulas et al., 2023).

FAO data collected globally since 1961 (FAOSTAT; https://www.fao.org/faostat/en/#data/QCL) show that Crop yields for maize, rice, wheat and soybean have increased as a result of enhanced irrigation, fertilization, and mechanization, as well as continuous selective breeding.

Although yields have doubled due to these methods, yield per hectare has plateaued recently, indicating that further investment in the current yield enhancement strategies may not result in higher output. There may be a chance to increase yield potential through photosynthesis.

One strategy to improve the efficiency with which intercepted radiation is turned into biomass is by increasing photosynthesis efficiency. Photosynthesis is a key mechanism that fixes carbon dioxide, generates oxygen, provides the energy that drives most life in the biosphere, regulates climate, and it underpins crop production. Additionally, improvements are required in plants’ ability to convert absorbed light energy into ATP and NADPH required to drive CO_2_ fixation by Rubisco and in minimizing carbon loss through processes such as photorespiration while protecting the system from photoinhibition (Garcia et al., 2023; Muhie, 2022).

Light is one of the most variable factors, forcing photosynthetic systems to adjust in order to maximize incoming energy and avoid photo inhibition (Salvatori et al., 2022). It is important to investigate how the plant canopy’s varying light availability temporal pattern affects physiological responses. Light-harvesting complexes may sustain photooxidative damage from incoming solar energy that surpasses saturation rates (Garcia et al., 2023).

The impact of the slow response of photosynthesis to fluctuating irradiance means that understanding the light adaptation mechanisms. These are crucial for improving photosynthesis and increasing overall crop yield by reducing limiting processes or selecting/ developing varieties well-adapted to fluctuating light environments Photosystem II is protected from damage through the photosystem II antenna system’s regulated thermal dissipation of absorbed light energy. This reversible photoprotective process reduces CO_2_ assimilation and photosystem II’s maximum quantum yield. This accounts for around a 15% reduction in daily canopy carbon uptake, with significantly higher values under stress (Long et al., 1994). Photosynthetic induction, or the increase in photosynthetic activity following light exposure following a dark-adapted state, can cause a leaf to take several minutes to reach optimal photosynthetic rates under specific conditions. Increasing the rate of induction may increase net photosynthetic performance in some scenarios (Murchie and Niyogi, 2011). This issue of slow induction is related to canopy structure, as lower canopy layers experience a greater fraction of their daily irradiance sum in the form of lightflecks.

The influence of fluctuating light on photosynthesis and overall carbon (C) uptake by plants is further complicated by the fact that it has been examined on different species under different growing conditions. Several leaf-scale studies have argued that light variations may reduce daily integral C assimilation (Morales and Kaiser, 2020), while others (Graham et al., 2017) have shown that specific fluctuation intervals and intensities may even boost photosynthesis. Light fluctuations cause plants to be in an unsteady state, and their capacity to adapt quickly to such variations is mostly determined by the effectiveness of antenna complexes (Kaiser et al., 2018a). The light-harvesting complex II (LHCII) is responsible for regulating energy distribution between PSI and PSII and the thermal dissipation of excess energy. However, crop yields have yet to benefit from this research, despite several tantalizing examples of genetic interventions in models (*Arabidopsis* and tobacco), including field experiments.

Temperature is a major meteorological factor influencing crop productivity and development. Temperature impacts enzyme activity within a leaf and causes changes in the developmental growth stage, which are closely linked to crop yield. Furthermore, the amount of water vapour in air at saturation grows exponentially with temperature, increasing the vapour pressure deficit (VPD) and driving more water loss from plants (Grossiord et al., 2020). As a result of these broad crop physiological reactions to temperature, any changes in long-term mean annual temperature occurrences are likely to have a large influence on crop productivity in the world’s key food and fuel-growing regions (Moore et al., 2021). Conversely, low temperatures can impede growth, leading to sink-limited photosynthesis or end-product photosynthesis (Harbinson and Yin, 2023). Low, non-freezing temperatures (between 5 and 15°C) have a deleterious impact on the growth and development of most plants. Plants are often exposed to low temperatures at some stage and have evolved methods to cope with such conditions. These adaptations may include changes in enzyme activity, osmolyte accumulation, changes to membrane fluidity (Upchurch, 2008), or specific modifications to photosynthesis and energy metabolism (Hüner et al., 2013). The net CO_2_ assimilation in C3 plants is primarily limited by Rubisco capacity at low CO_2_ levels. In contrast, at higher CO_2_, the limitation is shifted to ribulose bisphosphate (RuBP) regeneration capacity at suboptimal temperatures (Sage & Kubien, 2007).

Low-temperature stress can diminish stomatal conductance, light utilisation efficiency, and chlorophyll concentration in the leaf, ultimately lowering photosynthetic capacity (Li et al., 2015). Crop leaf photosynthesis is severely inhibited during cold stress, and requires a recovery period under non-stress conditions (Dikšaitytė et al., 2019). The recovery rate of photosynthetic capacity following low-temperature stress is determined by the extent of damage sustained during the stress and the climate condition after the stress is removed (Whaley et al., 2004). Despite temperature increases that follow stress, the leaf’s stomatal conductance and photosynthetic capacity take several days to recover. Temperature influences potential photosynthetic capacity at the leaf level, and additionally biomass output, by influencing the effective photosynthetic duration and area at the canopy scale (Xiao et al., 2022). The effect of low temperature on plants is determined not only by the temperature but also by other environmental factors to which the plant is simultaneously exposed (Waraich et al., 2012), particularly light-intensity (Franklin et al., 2014) and developmental stage at cold exposure (Prinzenberg et al., 2020).

Tomato, a widely grown species that originated in Central and South America, is one of the most economically important crops and has been in Europe since the 16th century. According to FAO data, it is the second most important vegetable crop, following potato. In 2021, the total area and production of tomatoes were 5.17 × 10^6^ ha and 1.89 × 10^8^ tonnes, respectively (Liu et al., 202). According to USDA statistics, it is one of the lowest-calorie vegetables and is high in antioxidants, dietary fiber, minerals, and vitamins. Researchers have worked hard to reintroduce lost features from wild relatives into commercial cultivars (Ye and Fan, 2021). The availability of the genome sequence (Tomato Genome Consortium, 2012) and the comparatively small genome size of 900Mb makes tomato an attractive crop for gene characterization, cytogenetics, molecular genetics, and molecular biology. Several traits related to its horticultural value have been extensively studied, such as size, pigment content, and flavor substances, with a focus on the metabolic and regulatory networks (Tieman et al., 2017). These qualities make tomato has been chosen as a model species for research.

In this study, four different S.*lycopersicum* genotypes were used to explore the short-term response of photosynthesis to low temperatures and fluctuating light. We look at the response of photosynthesis measured using modulated chlorophyll fluorescence imaging, to understand how the quantum efficiency of electron transfer through PSII (Φ_PSII_) changes to a step increase and step decrease in irradiance. We also look at how these responses are affected by prolonged cold exposure and subsequent recovery.

## MATERIALS AND METHODS

### Plant material and growing conditions

In this study, four diverse genotypes; Nagcarlan (P1), North Carolina heatset-1 (NCHS-1) (P2), Delfo parent 1(P3) and Delfo parent 2 (P4) and a control; *S. lycopersicum* (cv. Moneymaker) (MM) were cultivated under controlled climate conditions at the Netherlands Plant Eco-phenotyping Centre (NPEC, Wageningen). This growth room is equipped with PlantScreen^TM^ Robotic XYZ System which is a robotic arm designed for plant cultivation and growth monitoring. It moves laterally, height-wise, and vertically, delivering imaging sensors and irrigation units to plants. The system operates in a sensor-to-plant concept, allowing user-defined protocols (https://plantphenotyping.com/products/plantscreen-robotic-xyz-system/#details). The growth chamber is equipped with LED illumination, including natural daylight simulations, imaging equipment for the visible light spectrum (RGB), and chlorophyll fluorescence imaging (https://growth-chambers.com/products/growth-capsule-gc/#details). The cultivar Nagcarlan (LA2661) is heat tolerant, producing more pollen under long-term moderate heat. NCHS-1 (LA3847) is a country race with medium heat tolerance that produces a high pollen count (Xu et al., 2017). F1 hybrids Delfo Parents 1 and 2 (BASF, Nunhems, The Netherlands) are pest-resistant.

Seeds were germinated on 4cm diameter rockwool Plugs (Grodan, Roermond, the Netherlands) and kept in the dark at 21°C for three days. Seedlings were then transferred to rockwool blocks (10×10×6.5cm), 3 days after sowing (DAS). An in-house nutrient solution (Tomato 2.0)was used to irrigate the plants twice a week. All genotypes were grown in 3 replicates with a day length of 16 hours (from 7:00h to 23:00h), a light intensity of 400 µmol m^−2^ s^−1^, day/night temperatures of 21 / 18°C, and relative humidity of 75%. The day/night temperatures were lowered to a day temperature of 14°C and a night temperature of 11°C 19 days after sowing (19 DAS). The plants were grown in the cold for 6 days before the temperatures were elevated to 21°C and 18°C for day and night on day 25 after sowing.

### Phenotyping for photosynthetic responses

For the control experiment (13 DAS – 18 DAS), chlorophyll fluorescence was measured with an automated camera system that moved over the plants. Pre-dawn Fv/Fm measurements using saturating light flashes (8000 μmol m^−2^s^−1^, for 800ms) were taken before the lights went on at 5:30. Φ_PSII_ was measured at growth light irradiance twice a day, at 8:00h (Dawn) and 21:00h (Dusk). Responses of Φ_PSII_ to step decrease (200 µmol m^−2^ s^−1^) and step increase (600 µmol m^−2^ s^−1^) in irradiance were measured using a series of saturating light flashes at 12:00h. The Robotic XYZ System was used to equilibrate the plants for 2 minutes at 400 μmol m^−2^ s^−1^, and Φ_PSII_ was monitored every 30 seconds. The light intensity in the camera head was dropped to 200 µmol m^−2^ s^−1^ immediately after the last Φ_PSII_ measurement, and measurements of Φ_PSII_ were then taken for 15 minutes (every 5s for 25s, every 10s for 40s, every 30s for 1min, and then every 120s). After 15 minutes, the light intensity was increased to 600 µmol m^−2^ s^−1^ and the Φ_PSII_ measurements were repeated for 15 minutes with the same measurements as step decrease. The same protocol was repeated for all genotypes under cold (19 DAS-24 DAS) and recovery (25 DAS – 30 DAS) conditions. To quantify the responses of PhiPSII to step decrease and step increase in irradiance the average kinetic responses of each genotype were fit with the following equations.

### Calculations

#### Photosynthetic parameters

Maximum quantum efficiency of PSII,

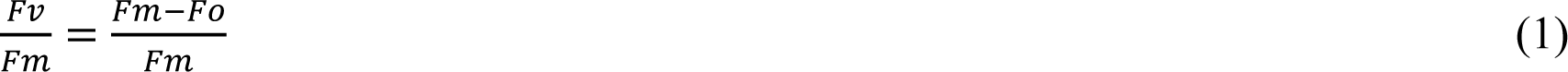

Where, Fm is maximum fluorescence when all reaction centers are closed, in the dark Fo is minimum fluorescence when all reaction centers are open, in the dark

Quantum efficiency of electron transport through PSII,

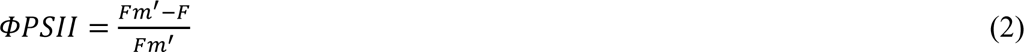

Where, Fm’ is maximum fluorescence when all reaction centers are closed, in actinic light F is steady-state fluorescence obtained, in actinic light

#### Fitting of kinetics

The curves were fit using the equation,

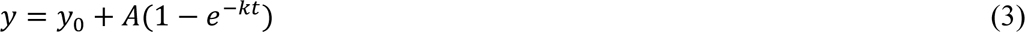

Where, y_0_ is the baseline value

A is the amplitude of response

k is the rate constant

## RESULTS

### Summary of growth conditions and phenotyping

The genotypes used for this study, Nagcarlan (P1), North Carolina heatset-1 (NCHS-1) (P2), ‘Delfo parent 1’(P3) and ‘Delfo parent 2’ (P4) and a control; *S. lycopersicum* (cv. Moneymaker) (MM), were grown for eighteen days at 21/18°C (day and night; ‘control’ treatment). The genotypes were then subjected to a cold treatment (14/11°C, day and night; ‘cold’ treatment) at the start of the day (00:00h) on the 19^th^ day after sowing (DAS). Furthermore, to understand whether the effects seen on photosynthesis were reversible in nature, the temperature was restored to the ‘control’ tomato growth temperatures (21/18°C, day and night; control treatment) on the 25^th^ DAS(00:00h).

### Influence of cold on the maximum quantum yield of PSII photochemistry, Fv/Fm

The maximum dark-adapted quantum efficiency of PSII photochemistry (Fv/Fm) was measured daily, 2.5 hours before the lights were switched on. The Fv/Fm was measured to check whether there is any photoinhibition, which can be caused by stress (Krause and Somersalo, 1989) or incomplete relaxation of photoprotective mechanisms of Photosystem II (PSII) (Demmig-Adams and Adams III, 2006). At the control temperature, the average Fv/Fm of all genotypes ranged from 0.76 to 0.81(16 DAS to 18 DAS; Fig 1), indicating the absence of stress or very little stress. P1 had consistently the most variable and lowest Fv/Fm (∼0.76) whilst P3 and P4 had consistently the least variable and highest Fv/Fm (∼0.81).

**Figure 1:**
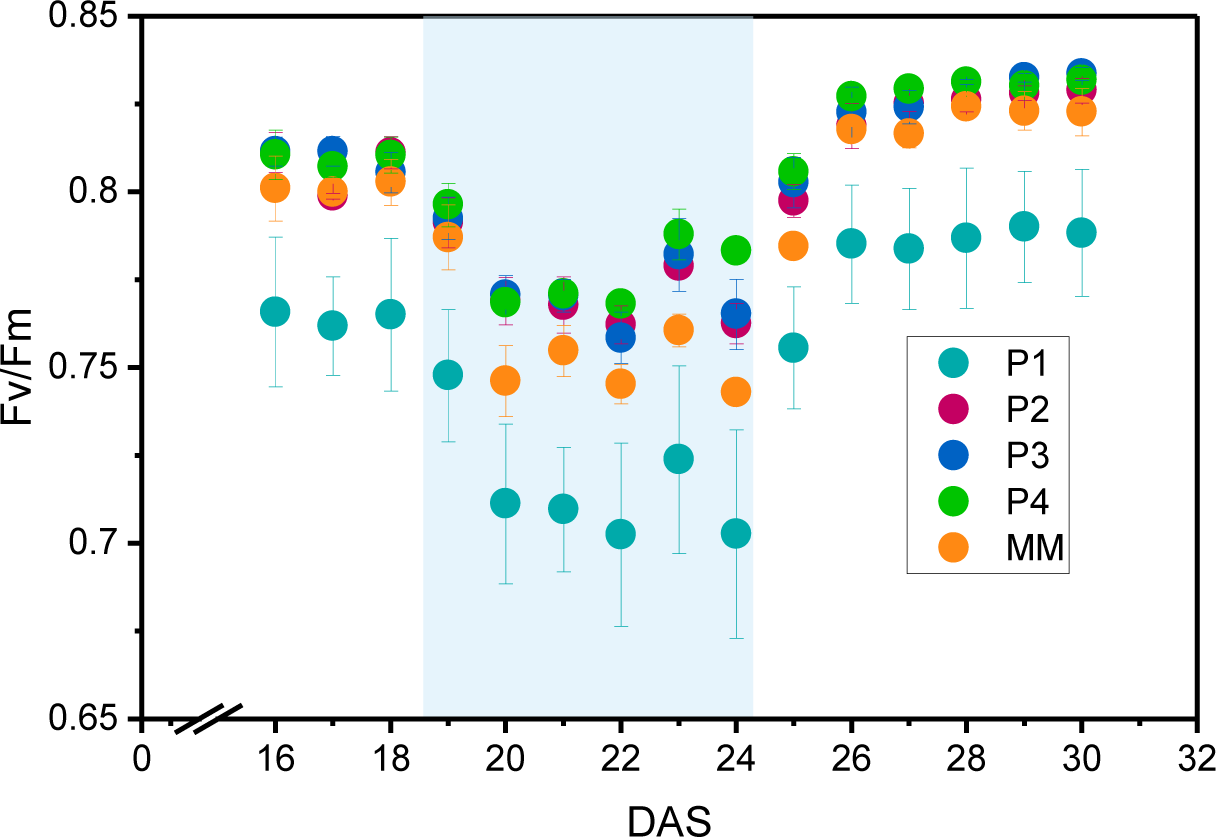
Scatter plot illustrating Maximum quantum efficiency, Fv/Fm, for all genotypes, measured each day after sowing (DAS). Fv/Fm measured 1h before the lights were switched on. The cold treatment started at 19 DAS and lasted till 24 DAS (including). The days with cold exposure are indicated in light blue. Each point indicates the mean (n = 3) ΦPSII and the error bar indicates standard error.

After the temperature was lowered to 14^°^C(19 DAS), the average Fv/Fm of all genotypes (measured after 5.5h in cold) dropped slightly (Fig 1; 19 DAS). The average Fv/Fm of P1, P2 and MM dropped by 2.2% (0.76 to 0.74, 0.81 to 0.79 and 0.80 to 0.78 respectively)while P3 and P4 saw a drop in Fv/Fm of 1.6% (0.8 to 0.79 and 0.81 to 0.8 respectively) compared with last day of control (18 DAS). With prolonged exposure to the cold, the Fv/Fm of all genotypes dropped further and reached a minimum at 22 DAS. However, the effect of cold on the Fv/Fm was more severe for all genotypes compared to the start of the cold treatment (19 DAS). This is apparent from the degree to which the Fv/Fm dropped at the end of cold exposure (Fig 1; 24 DAS) compared to the last day of control. The Fv/Fm for P1 dropped the most, from 0.76 to 0.70 (∼8%), followed by MM, which had a drop in Fv/Fm of 7.4% (0.8 to 0.74). P2 and P3 experienced a drop in Fv/Fm of ∼5% (0.81 to 0.76,0.8 to 0.76 respectively) while P4 showed the lowest drop of 3.3% (0.81 to 78). This suggests that the Fv/Fm of P4 is the least sensitive to cold while the Fv/Fm of P1 is quite sensitive to cold. When the temperature was restored to 21°C (25 DAS), the Fv/Fm of all genotypes increased. Most notable was the recovery Fv/Fm of P1, which increased the most, by 7.5%, from 0.70 to 0.75, in 5.5h at 21°C. While the Fv/Fm of MM, P2 and P3 increased, around 5.5% for MM (0.74 to 0.78) and by 4.8% for P2 and P3 (0.76 to 0.80), P4 saw the least recovery with only 2.8% (0.78 to 0.80) increase in Fv/Fm (Fig 1; 25 DAS). The low recovery of the Fv/Fm of P4 is unsurprising, as the Fv/Fm of P4 was high during the control conditions and dropped the least under prolonged cold exposure, hence the degree of possible recovery is also lower. Additionally, the degree of recovery of all genotypes were higher than the degree to which the Fv/Fm dropped. The Fv/Fm of all genotypes reached a maximum under ‘recovery’ conditions at 28 DAS. At the end of the recovery period (Fig 1; 30 DAS), P2,P3 and P4 have similar Fv/Fm of 0.83, whilst MM and P1 had an Fv/Fm of 0.82 and 0.78 respectively, indicating that the effect of cold on the Fv/Fm is completely reversible.

### Response of the quantum efficiency of PSII photochemistry (Φ_PSII_) to cold exposure and recovery

In order to see how the effect of cold influences photosynthesis and to see whether these changes, like the Fv/Fm, were reversible, the steady-state quantum efficiency of electron transport through photosystem II (Φ_PSII_) was measured. The Φ_PSII_ was measured at 400 μmol m^−2^s^−1^ (growth light conditions; see methods) 1h after the light is switched on (referred to as ‘dawn Φ_PSII_’) and Φ_PSII_ measured 1h before the light was switched off (referred to as ‘dusk Φ_PSII_’). The dawn Φ_PSII_ and the dusk Φ_PSII_ was taken to monitor how Φ_PSII_ changes at the end of the day compared to the start of the day. Under control temperatures (21°C), P1 had the lowest Φ_PSII_ (0.63) whilst all other genotypes had similar quantum efficiencies (0.67) (Fig 2 A,B 16-18DAS).

**Figure 2:**
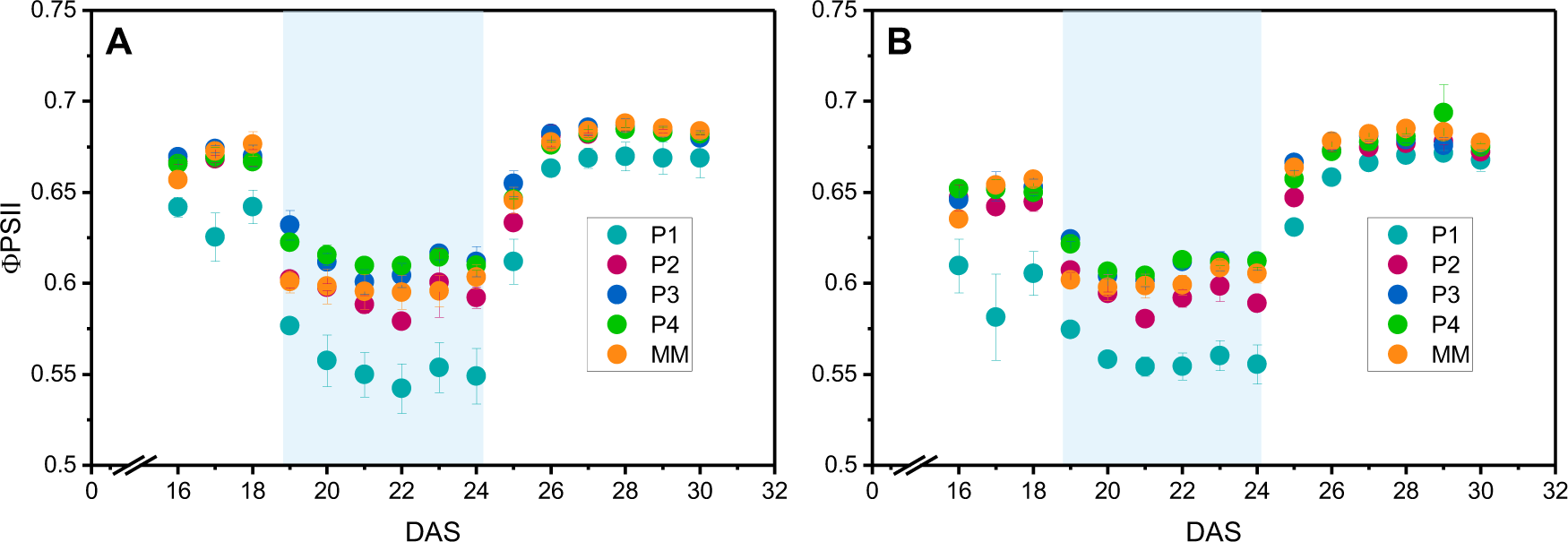
Scatter plot illustrating Quantum efficiency, ΦPSII, for all genotypes, measured each day after sowing (DAS). (**A)** ΦPSII measured 1h after the lights were turned on (Dawn), measured at 400 μmol m^−2^ s^−1^. (**B)** ΦPSII measured 1h before the lights were switched off (Dusk), measured at 400 μmol m^−2^ s^−1^. The cold treatment started at 19 DAS and lasted till 24 DAS (including). The days with cold exposure is indicated in light blue. Each point indicates the mean (n = 3) ΦPSII and the error bar indicated standard error.

After subjecting all genotypes to cold (19 DAS) both the dawn and the dusk Φ_PSII_ dropped, and kept dropping till they reached a minimum at 22 DAS (Fig 2 A, B). At 19 DAS, the average dawn Φ_PSII_ of MM dropped the most from 0.68 to 0.6 (11.1%), followed by P1 and P2 (10%; 0.66 to 0.6 and 0.64 to 0.57, respectively), of which P1 had the lowest Φ_PSII_ (0.57) amongst all genotypes. P3 and P4 on the other hand, had significantly lower drops in Φ_PSII_, only dropping 5.6%(0.67 to 0.63) and 6.6% (0.66 to 0.62), respectively (Fig 2; 19 DAS). For the Φ_PSII_ observed at ‘dusk’ the trend remained the same as the dawn measurements, with MM and P1 having the largest drop, compared to the last day of control, followed by P2 and the lowest drops for P3 and P4. When comparing the dawn Φ_PSII_ on the last day of the cold treatment (24 DAS; Fig 2A) to the dawn Φ_PSII_ on the last day of control (18 DAS; Fig 2A) P1 experienced a considerable loss of 14% (0.64 to 0.54)while all the other genotypes experienced ∼9-10% reduction in Φ_PSII_. The reduction in dusk Φ_PSII_ is less severe when comparing 24 DAS to 18 DAS, where all the genotypes experience 6-8% drop in Φ_PSII_.

Following the restoration of the temperature to 21°C (25 DAS) the dawn and dusk Φ_PSII_ of all genotypes recovered to pre-cold values in two days (26 DAS; Fig 2). After 5.5h at 21°C (25 DAS), all genotypes showed a considerable increase in the dawn Φ_PSII_, where P1 recovered from 0.54 to 0.61 (11%) and all the other genotypes recovered around (8%; 25 DAS Fig 2A). A similar extent of recovery was observed for the dusk Φ_PSII_, where P1 had the highest recovery of 13% (0.55 to 0.63) and while the other genotypes experienced an 8-9% recovery (25 DAS; Fig 2B). Notably, after recovery, all genotypes had a Φ_PSII_ that was higher than those observed under control conditions. This was especially true for the dusk Φ_PSII_ where all genotypes had an average Φ_PSII_ of 0.67(30 DAS; Fig 2B), whereas on the last day of control, the average Φ_PSII_ of all genotypes except P1(0.6) was 0.65.

### Short-term response of photosynthesis to a step increase and step decrease in irradiance

Photosynthesis responds to changes in irradiance in the short term. The response of photosynthesis to increases and decreases in irradiance depends on the responses of different processes so it is necessary to investigate responses to both. (Kaiser et al., 2018b). Hence, to investigate how the photosynthesis of each genotype responds differently to short-term changes in irradiance all genotypes were subjected to a step increase and step decrease in irradiance. The initial and final irradiance values for both the step increase and step decrease were selected based on the irradiances that span a large change in CO_2_ assimilation of the control genotype, MM, inferred from a light response curve (Supplementary Fig S1). For the step decrease, a transition from 400 μmol m^−2^ s^−1^ to 200 μmol m^−2^ s^−1^ was selected while for step increase a transition from 200 μmol m^−2^ s^−1^ to 600 μmol m^−2^ s^−1^ was adopted. The effect of these short-term changes in irradiance on photosynthesis was carried out by monitoring Φ_PSII_. Φ_PSII_ was used to monitor photosynthesis for two reasons: (1) Φ_PSII_ is a major determinant of linear electron transport, and hence, photosynthetic Light Use Efficiency (LUE); Improving LUE is integral in improving photosynthesis (2) Φ_PSII_ can be measured by imaging chlorophyll fluorescence with the application of saturating light flashes, allowing for a non-contact, non-destructive measurement of fast changes in photosynthesis with good spatial and temporal resolution.

Figure 3 shows the changes in Φ_PSII_ measured with time, monitored with the application of saturating light pulses, during transitions from a lower to a higher irradiance and vice-versa. During the transition from 400 μmol m^−2^ s^−1^ to a lower light intensity of 200 μmol m^−2^ s^−1^ Φ_PSII_ of all genotypes increased rapidly and slowed down to eventually reach a new steady state (Fig 3. A). Whilst, during the transition from 200 μmol m^−2^ s^−1^ to 600 μmol m^−2^ s^−1^ Φ_PSII_ dropped immediately and slowly recovered to a new steady state (Fig 3B).

**Figure 3:**
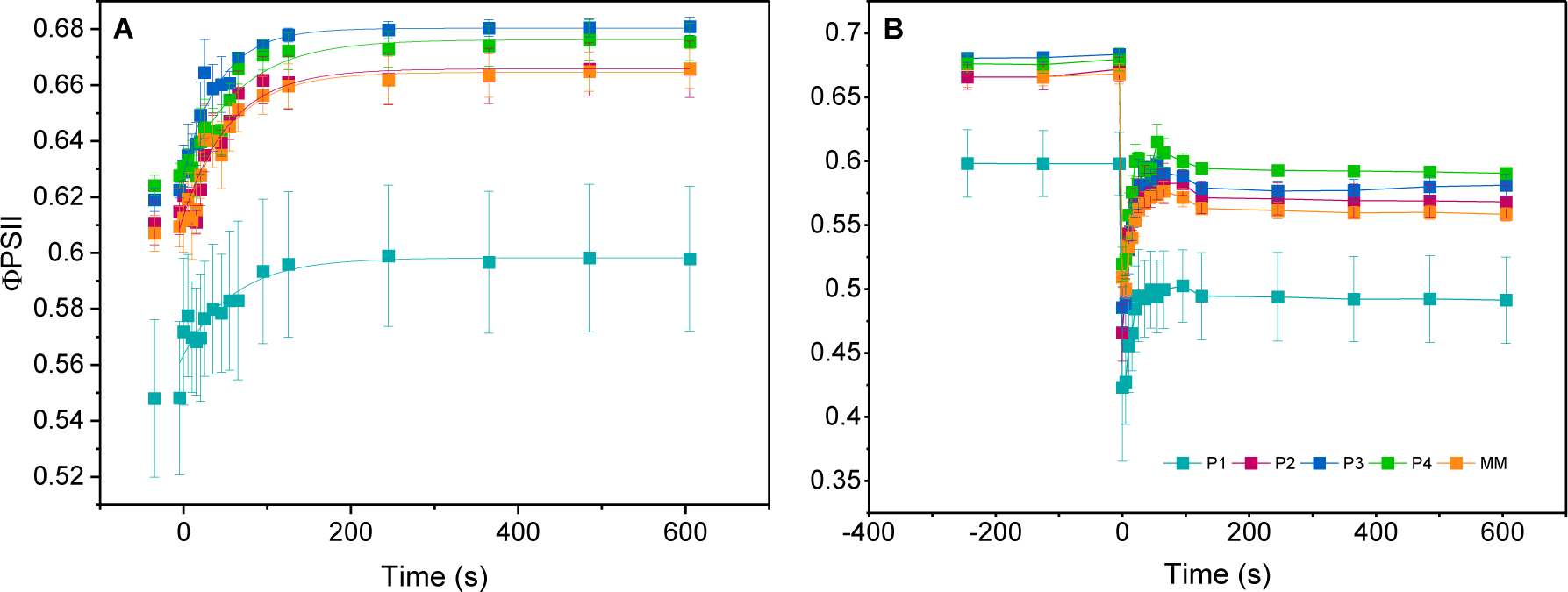
Scatter plot illustrating kinetics of ΦPSII for all genotypes (12 DAS) following a step change in irradiance at 21°C. **(A)** Illustrates the kinetics for a step decrease in irradiance from 400 μmol m^−2^ s^−1^ to 200 μmol m^−2^ s^−1^. Solid lines indicate the fit curve. **(B)** Illustrates the kinetics for a step increase in irradiance from 200 μmol m^−2^ s^−1^ to 600 μmol m^−2^ s^−1^. Solid lines are the linear connections between each point. Each point shows a mean value (n = 3) and error bars indicate standard deviation

To compare how fast photosynthesis of each genotype responds to the short-term change in irradiance the time course change in Φ_PSII_ was fit with a mono-exponential function (equation 3; see calculations), and the consequent fit parameters are summarized in Table 1. Based on the first order rate constant (k), P2 seem to respond the fastest during the transition from high to low irradiance while all the other genotypes seem to respond equally fast (Table 1). On the other hand, during the step increase in light intensity, most genotypes showed similar rates of response, except for MM, which had the lowest rate constant (Table 1). When compared to the other genotypes P1 had the lowest steady-state Φ_PSII_ (0.59) after a step decrease in irradiance. Similarly, P1 also had the lowest steady-state Φ_PSII_ after the step increase (0.49) in irradiance.

**Table 1:** Summary of fit parameters for both step increase and step decrease to irradiance at 21°C. The fit parameters are obtained from equation 3 (refer Calculations).The fit parameters are rate constants (k) and steady state reached after the step increase or step decrease (SS). The standard error (±SE) of the fits (n=3) is provided. Statistical test of Welch’s t-test was carried out (Supplementary Fig S2-5).*Statistically different from other values in the same column (p < 0.05).

| Genotype | Step Decrease |  | Step Increase |  |
| --- | --- | --- | --- | --- |
| | k (s<sup>-1</sup>) | SS ($\Phi_{PSII}$) | k (s<sup>-1</sup>) | SS ($\Phi_{PSII}$) |
| P1 | 0.018 ± 0.002 | 0.59 ± 0.002* | 0.10 ± 0.01 | 0.49 ± 0.02* |
| P2 | 0.031 ± 0.005* | 0.66 ± 0.004 | 0.10 ± 0.019 | 0.58 ± 0.01 |
| P3 | 0.021 ± 0.005 | 0.68 ± 0.004 | 0.09 ± 0.021 | 0.59 ± 0.02 |
| P4 | 0.023 ± 0.004 | 0.66 ± 0.005 | 0.11 ± 0.02 | 0.59 ± 0.006 |
| MM | 0.020 ± 0.004 | 0.66 ± 0.005 | 0.06 ± 0.01* | 0.57 ± 0.008 |

### Short-term response of photosynthesis to a step increase and step decrease in irradiance under low temperature conditions

The temperature at which the plants were grown at was dropped by 7°C, from 21°C to 14°C (00:00; 19 DAS), to understand how the response of photosynthesis to both a step increase and step decrease in irradiance changes with cold. Identical to what was carried out under control conditions, the change in Φ_PSII_ during the step changes were monitored. Figure 4 shows the kinetics of all genotypes to a step decrease (Fig 4A) and step increase (Fig 4B) in irradiance and the parameters obtained from the fit to Equation 3 are summarized in Table 2. Identical to what was observed under control conditions, during step decrease in irradiance, Φ_PSII_ of all genotypes increased rapidly and slowed down to eventually reach a new steady state (Fig 4A). Whilst during step increase in irradiance, the Φ_PSII_ of all genotypes dropped immediately and slowly recovered to reach a new steady state (Fig 4B). Additionally, unlike the pattern observed under control conditions, for both the step increase and step decrease to irradiance, the relative variation in average Φ_PSII_ values are larger under cold conditions (Fig 4).

**Figure 4:**
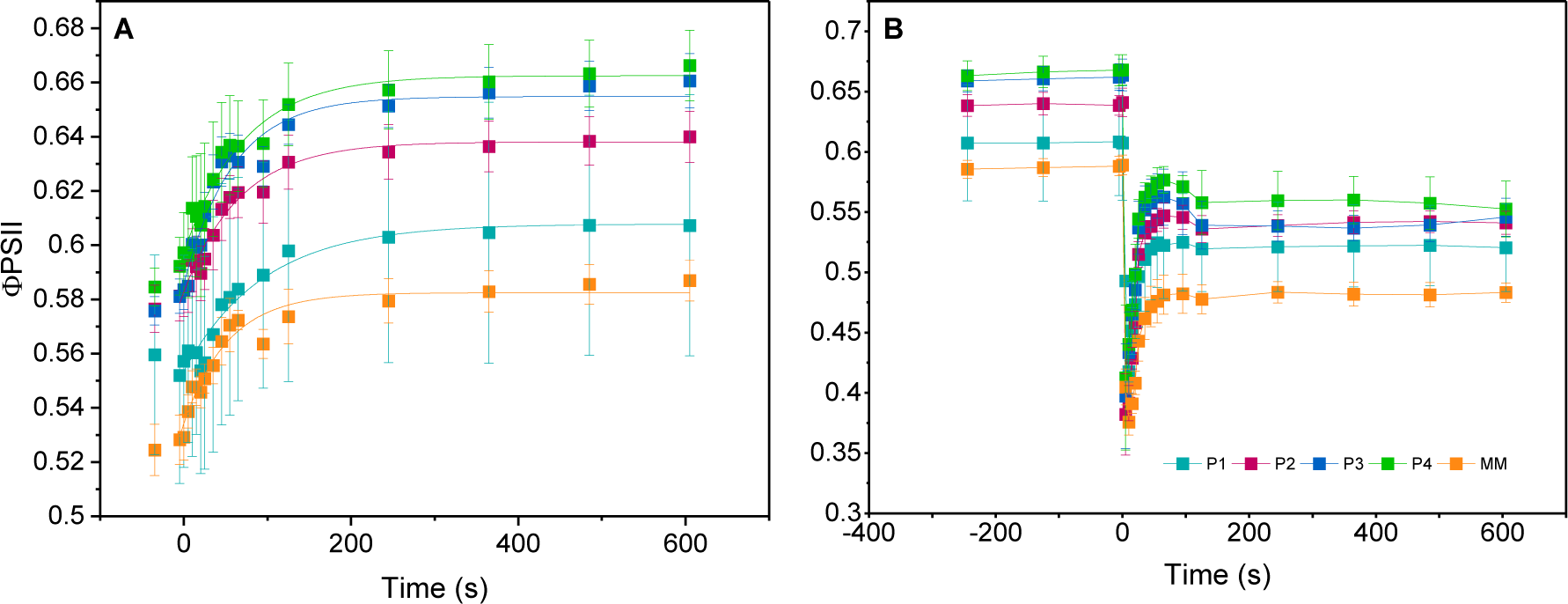
Scatter plot illustrating kinetics of, ΦPSII for all genotypes (19 DAS) following a step change in irradiance at 14°C. **(A)** Illustrates the kinetics for a step decrease in irradiance from 400 μmol m^−2^ s^−1^ to 200 μmol m^−2^ s^−1^. Solid lines indicate the fit curve. **(B)** Illustrates the kinetics for a step increase in irradiance from 200 μmol m^−2^ s^−1^ to 600 μmol m^−2^ s^−1^. Solid lines do not indicate the fit curve. Each point shows a mean value (n = 3) and error bars indicate standard deviation.

**Table 2:**
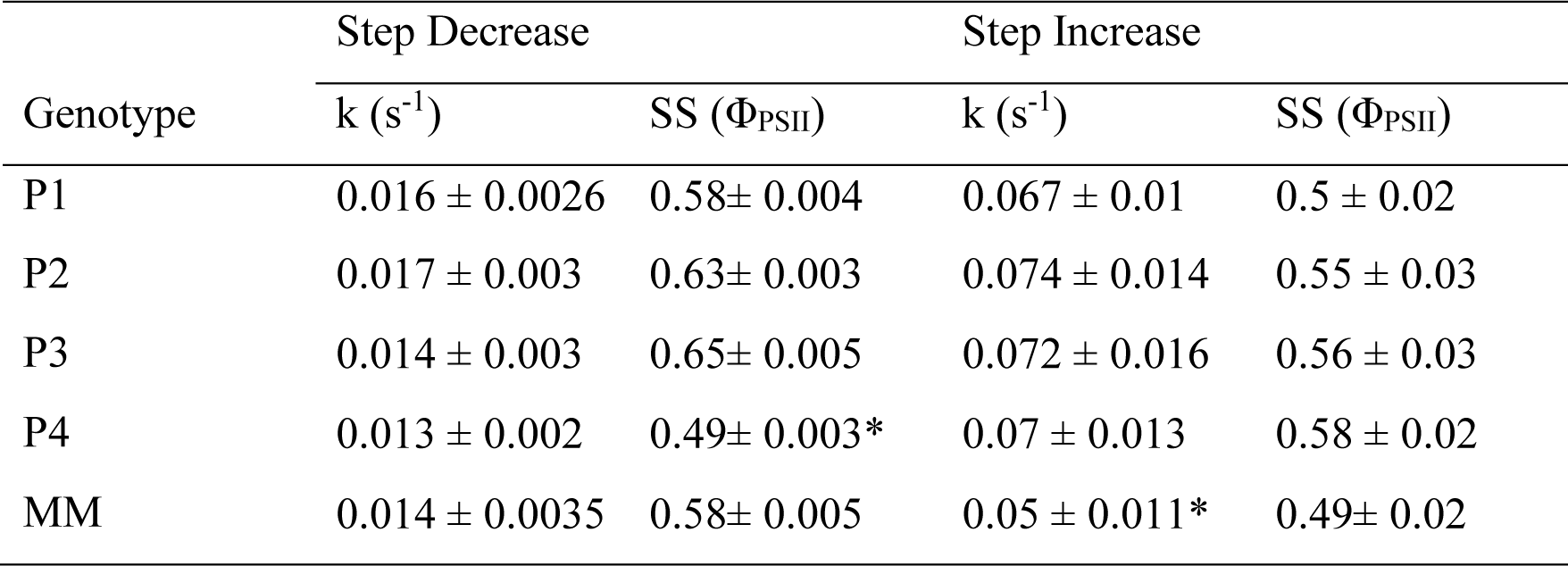
Summary of fit parameters for both step increase and step decrease to irradiance at 14°C. The fit parameters are obtained from Equation 3 (refer Calculations). The fit parameters are rate constants (k) and steady state reached after the step increase or step decrease (SS). The standard error (±SE) of the fits (n=3) is provided. Statistical test of Welch’s t-test was carried out (Supplementary Fig S2-5).*Statistically different from other values in the same column (p < 0.05).

Comparing the fit parameters, the rate constants of all genotypes during step decrease in irradiance drops slightly. However, the rate constants of all genotypes of the response to step increase in irradiance dropped significantly (Table 2), when compared to the control condition (Table 1), of which the rate constant of MM was the lowest (Table 2). When comparing steady-state Φ_PSII_ after a step decrease in irradiance to those under control conditions, the Φ_PSII_ of P4 and MM (lowest Φ_PSII_; 0.49) dropped the most (Table 2). However, of all the genotypes, the Φ_PSII_ after step increase in irradiance of MM dropped the most.

### Change in kinetics of short-term responses of photosynthesis with cold stress and recovery

The rate constants of the response of Φ_PSII_ and the steady-state Φ_PSII_ of all genotypes were influenced by the exposure to cold (19 DAS). Response of Φ_PSII_ to a step increase and step decrease to irradiance was measured at the end of the cold exposure (24 DAS). This was done to understand how prolonged cold exposure will affect the rate constants and steady-state Φ_PSII_. Additionally, to investigate whether the effect of cold on the rate constants and steady-state Φ_PSII_ were reversible the response of Φ_PSII_ to the step changes in irradiance was measured at the start of ‘recovery’ (25 DAS) and at the end of the ‘recovery’ phase.

Fig. 5 highlights how the rate constant of photosynthetic response (Φ_PSII_) during both step increase and step decrease in irradiance changes with changes in temperature. Notably, the rate constants obtained from response to step decrease in irradiance of most genotypes remain relatively unchanged with drop in temperature and recovery (Fig 5; left). For genotypes P2 (Fig 5C; cold) and P4 (Fig 5G; cold), where the rate constants drop with cold, the rate constants recover back after the recovery period. On the other hand, for the response to step increase in irradiance, all genotypes except P2 and P4(Fig 5 D,H) showed a marked decrease in the rate constant under cold conditions (Fig 5; right; cold). P1 showed only a slight decrease in the rate constants (Fig. 5 B), while the effect was much more prominent P3 and MM (Fig 5 F,J). The rate constants for all genotypes remained the same with prolonged exposure to cold (Fig 5 B,D,F,H,J; cold_acc). Surprisingly, the rate constants, during the ‘recovery’ phase for P1 (Fig 5 B, Rec) and MM (Fig 5 J; Rec) were higher than during the control conditions. The rate constant of P3 on the other hand increased gradually to pre-cold levels (Fig 5 F; Rec and Rec_acc)

**Figure 5:**
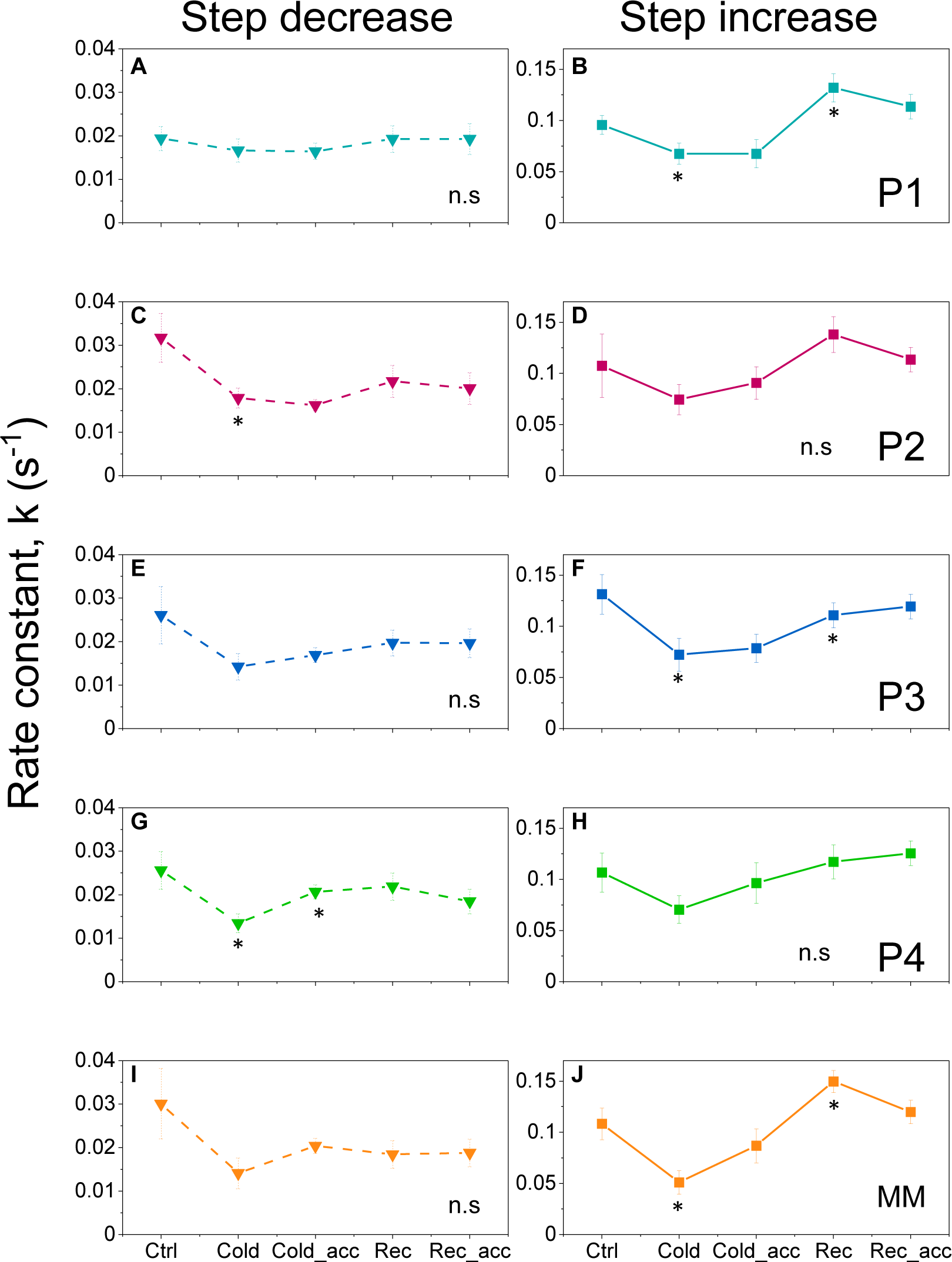
Line plot indicating the trend for kinetics to step change in irradiance during different conditions. Each point represents kinetics under different conditions as follows: Control (Ctrl; 18 DAS), Cold (19 DAS), Cold acclimation (Cold_acc; 24DAS), Recovery (Rec; 25DAS), acclimation to recovery conditions (Rec_acc; 30DAS). **(A, C, E, G**, and **I)** illustrate changes in kinetics to a step increase in irradiance from 200 μmol m^−2^ s^−1^ to 600 μmol m^−2^ s^−1^ for P1, P2, P3, P4 and MM respectively. **(B, D, F, H, and J)** illustrate changes in kinetics to a step decrease in irradiance from 400 μmol m^−2^ s^−1^ to 200 μmol m^−2^ s^−1^ for P1, P2, P3, P4 and MM respectively. Each point shows a mean value (n = 3) and error bars indicate standard error (±SE). Statistical test of Welch’s t-test was carried out (Supplementary II & IV).*Statistically different from the point to the left of it (p < 0.05). ‘n.s’ indicates no point is significantly different from each other

### Changes in steady-state Φ_PSII_ attained following step changes in irradiance under cold stress and recovery

While the kinetics (rate constants) are crucial in understanding how fast each genotype responds to each condition, the changes in the steady-state values of Φ_PSII_, after step increase or step decease in irradiance, are also crucial as they indicate the efficiency of linear electron transport, and by extension photosynthetic light use efficiency. Hence, the changes in the steady state attained following a step increase or step decrease in irradiance after cold exposure, subsequent cold acclimation and succeeding recovery were compared.

After the initial exposure to cold (19 DAS) the steady-state Φ_PSII_ after step decrease in irradiance for all genotypes except P1 drop (Fig 6 A). At the end of the prolonged exposure to cold (Fig 6 left; Cold_acc), the Φ_PSII_ of P2 and MM (Fig 6 C,I) remained unchanged while the Φ_PSII_ of P1 and P4 increased (Fig 6 A,G). P3 was the only genotype where the Φ_PSII_ decreased after prolonged cold exposure (Fig 6 E; Cold_acc). The Φ_PSII_ of all genotypes increased when the temperature was restored to 21°C (Fig 6 left; Rec). The Φ_PSII_ during the ‘recovery’ phase of all genotypes except P4, were higher than the Φ_PSII_ observed before exposure to cold. This pattern of increased Φ_PSII_ in recovery phase compared to the control conditions were also observed for the steady state Φ_PSII_ at 400 μmol m^−2^ s^−1^ (Fig 2).

**Figure 6:**
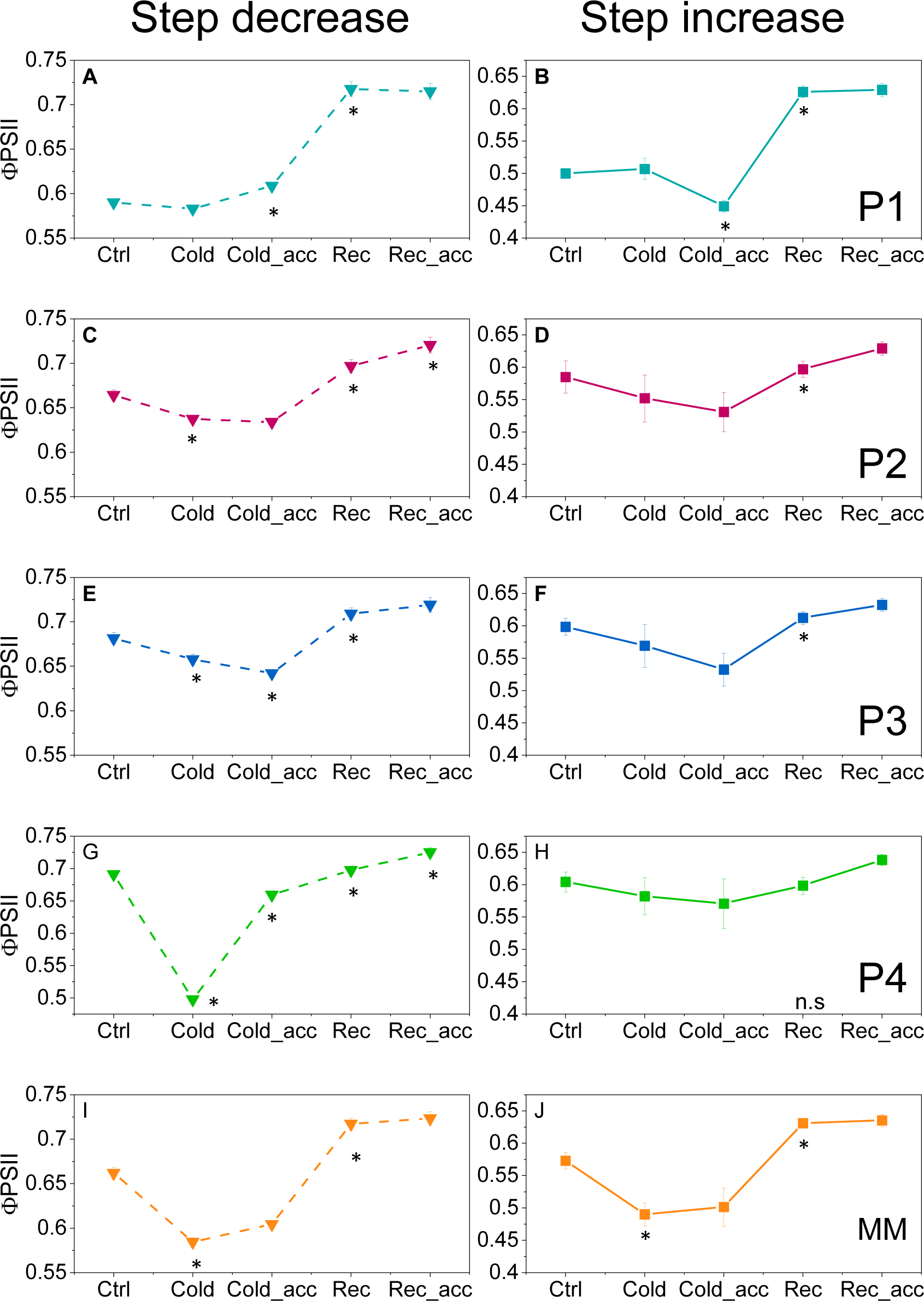
Line plot indicating the trend for steady-state Φ_PSII_ reached after a step change in irradiance during different conditions. Each point represents kinetics under different conditions as follows: Control (Ctrl; 18 DAS), Cold (19 DAS), Cold acclimation (Cold_acc; 24DAS), Recovery (Rec; 25DAS), acclimation to recovery conditions (Rec_acc; 30DAS). **(A, C, E, G**, and **I))** illustrate changes in kinetics to a step decrease in irradiance from 400 μmol m^−2^ s^−1^ to 200 μmol m^−2^ s^−1^ for P1, P2, P3, P4 and MM respectively. **(B, D, F, H, and J)** illustrate changes in kinetics to a step increase in irradiance from 200 μmol m^−2^ s^−1^ to 600 μmol m^−2^ s^−1^ for P1, P2, P3, P4 and MM respectively. Each point shows a mean value (n = 3) and error bars indicate standard error (±SE). Statistical test of Welch’s t-test was carried out (Supplementary III & V).*Statistically different from the point to the left of it (p < 0.05). ‘n.s’ indicates no point is significantly different from each other

The effect of cold on the steady-state Φ_PSII_ after step increase in irradiance was different to that observed during step decrease in irradiance. Only MM (Fig 6 J) had lower Φ_PSII_ than those observed during the control conditions. At the end of the cold treatment (24 DAS) the Φ_PSII_ of P1 dropped significantly (Fig 6 B; Cold_acc) while the Φ_PSII_ of all the other genotypes had the same Φ_PSII_ as the start of the cold. Much like steady state Φ_PSII_ after step decrease and Φ_PSII_ at growth irradiance, during the ‘recovery’ phase the Φ_PSII_ was higher than the control conditions. P4 was the only exception, wherein the Φ_PSII_ remains constant throughout the changes in temperature (Fig 6 H).

## Discussion

Acclimation of photosynthesis to fluctuating light and limitations to CO2 assimilation during transients has been studied extensively (eg. Kaiser et al., 2018a; Kaiser et al., 2020; Sassenrath-Cole and Pearcy, 1992; Sassenrath-Cole and Pearcy, 1994; Sassenrath-Cole et al., 1994). While some of the aforementioned studies (Sassenrath-Cole and Pearcy, 1992; Sassenrath-Cole and Pearcy, 1994; Sassenrath-Cole et al., 1994) look at the constraints during photosynthetic induction from darkness, none focus on the transient responses to both increasing and decreasing irradiances. Similarly, the effect of cold on photosynthesis for tomato plants has been studied (Brüggemann et al., 1996; Brüggemann et al., 1992; Martin and Ort, 1982; Martin and Ort, 1985; Martin et al., 1981; Sassenrath et al., 1990) but, the combined effect of cold and fluctuating light on photosynthesis has not received much attention. Here, we explored the effect of short-term fluctuating light (both increase and decrease of irradiance) and cold on photosynthesis in multiple genotypes of tomato. We observed a significant genotype-to-genotype variation in the photosynthetic responses both to short-term fluctuations and changes in temperature.

A decrease in Fv/Fm has been commonly used as a measure of plants’ response to stress. This is especially true in cold environments, (Krause and Somersalo, 1989; Ogaya et al., 2011; Rizza et al., 2001). We observed a decrease in Fv/Fm when the temperature was lowered to 14°C, however, the Fv/Fm recovered when temperatures were restored to 21°C (growth temperature), indicating that the effect of cold was largely reversible. The Fv/Fm of P1 was quite sensitive to cold, which was apparent from the largest drop in Fv/Fm in just 5.5h of cold compared to the other genotypes. Interestingly, while the Fv/Fm of P1 dropped significantly with 5.5h at 14°C, the Fv/Fm of P1 also recovered the most at 5.5h at 21°C. Additionally, the Fv/Fm of P4 showed the least change with prolonged exposure to cold only dropping by 3.3% to 0.78 whereas the other genotypes experienced a relatively large decrease in Fv/Fm.

With changes in temperature, the Φ_PSII_ of all genotypes also showed trends that were similar to those observed for Fv/Fm. This suggests that Φ_PSII_ could also be a useful indicator of the effect of cold on photosynthesis with the advantage of not requiring long dark adaptation. The dusk Φ_PSII_ were slightly lower than the dawn Φ_PSII_ (Fig 2), potentially due to the effect of the light treatment, circadian rhythm or other diurnal processes (Lanoue et al., 2018; Matthews et al., 2018). The effect of cold on both dawn and dusk measurements were quite similar, evidenced by the similar degrees of change in Φ_PSII_ observed for all genotypes. Similar to that observed for Fv/Fm, P1 had consistently low Φ_PSII_ in all temperature conditions. Also similar to the pattern observed with changes in Fv/Fm to temperature, the dawn and dusk Φ_PSII_ of P1 dropped the most during the ‘cold’ phase and also recovered the most during the ‘recovery’ phase. Additionally, MM experienced a huge drop in Φ_PSII_ after the onset of cold temperatures, but the Φ_PSII_ did not drop further like that observed for the other genotypes. Comparing the patterns in both Fv/Fm and Φ_PSII_, it can be inferred that both MM and P1 are especially sensitive to cold.

The results of our investigation have indicated strongly that the factors affecting steady-state properties of photosynthesis are different to those regulating the response of photosynthesis to a changing irradiance. The genotypes that responded fast or slow to changes in irradiances did not necessarily have highest or lowest Φ_PSII_ attained following those changes in irradiances. Moreover, the genotype that responded the slowest during step increase (MM), was not the slowest to respond to step decrease in irradiance. This agrees with the suggestion that processes that limit photosynthesis during an increase in irradiance are different from those limiting photosynthesis during a decrease in irradiance (Kaiser et al., 2018b; Kaiser et al., 2015). Studies on photosynthetic induction and relaxation have provided insight into the activation and deactivation of numerous processes that regulate photosynthesis (Kaiser et al., 2018a; Kaiser et al., 2020; Sassenrath-Cole and Pearcy, 1992; Sassenrath-Cole and Pearcy, 1994; Sassenrath-Cole et al., 1994); reviews (Kaiser et al., 2018b; Kaiser et al., 2015). However, the implication that different factors affect the kinetics and steady-state response of photosynthesis to fluctuating has not been reported, especially the consideration that these constraints might be different for different genotypes. This distinction can provide a better approach in understanding which processes limit photosynthesis under fluctuating light and specifically the genetic factors underlying these constraints.

Martin and Ort (1982) reported the decrease in quantum yield of PSII to chilling (1°C at 16h) measured via photoreduction of hexacyanoferrate(III) under growth light conditions and also observed the reduction in photosynthetic capacity, measured via CO_2_ saturated light response curve. This is similar to what we observed at 14°C, where the quantum yield of photosynthesis (Φ_PSII_) reduced both during growth light conditions (400 μmol m^−2^s^−1^) and after an increase and decrease in irradiance. The observed decrease in steady-state Φ_PSII_ following an increase or decrease in irradiance is similar to the reductions in CO_2_ assimilation observed by Powles et al. (1983) in *Phaseolus vulgaris* and Martin and Ort (1985) in *Solanum lycopersicum*, where they report a large drop in CO_2_ assimilation following the combined effect of temperature and high light. The rates constants of the changes in Φ_PSII_ to step increase and step decrease in irradiance were also affected by cold exposure. Powles et al. (1983) observed changes in CO_2_ assimilation and stomatal conductance with chilling temperatures (6°C) and Sassenrath et al. (1990) observed the loss of FBPase activity and reduction in RuBP concentrations under cold conditions (8°C). RuBP regeneration has been suggested to limit fast photosynthetic response in light flecks (Sassenrath-Cole and Pearcy, 1994; Sassenrath-Cole et al., 1994). Hence, a decrease in FBPase activity and reduction in stomatal conductance could potentially explain the changes in rate constants for the response to a step increase in irradiance observed under cold. However, Martin et al. (1981), however, observed a reduction in photosynthesis for chilled leaves (1°C) acclimated to CO_2_ saturated conditions (1500 ppm). Assuming that photosynthesis is not limited by chloroplast CO_2_ (C_c_) concentrations at 1500 ppm CO_2_, could suggest that metabolic limitations and not stomatal limitations limit photosynthetic response to short-term fluctuations in irradiance during cold exposure. The most notable observation was that the rate constant during a step decrease in irradiance was less affected by cold, especially P1, which showed no significant change in rate constants with temperature. This relative insensitivity of rates of response of photosynthesis to step decrease in irradiance has not been reported previously. Barrero-Gil et al. (2016) have also reported changes in the metabolome of cold-adapted and cold-exposed leaves (10°C). This could also explain the variation in the observed photosynthetic responses under prolonged exposure to cold and recovery, especially the higher steady-state Φ_PSII_ observed after recovery from cold for some genotypes.

Comparing genotypes, we observed that the rate constant for the response of P1 to decrease in irradiance was unchanged while there was relatively less change in the rate of response to an increase in irradiance with cold and even prolonged cold exposure. Similarly, P2, P3 and P4 had relatively unchanged Φ_PSII_ throughout the cold and recovery conditions in response to step increase and step decrease in irradiances. The genetic factors that influence the observed plasticity of the rate constants demonstrated by P1 and the relatively unchanged Φ_PSII_ of P2, P3 and P4 could serve to be indicators of potential markers for desirable traits to improve photosynthesis under fluctuating light and cold. This can result in the LUE, influenced by the rate constants and the steady-state Φ_PSII_ after step changes to irradiance, decreasing with cold, at least under the limited range of temperatures employed here. Moreover, P1 and P2 are known heat-tolerant lines for pollen viability, albeit not investigated for heat tolerance for photosynthesis or vegetative growth (Villareal et al., 1977; Xu et al., 2017a; Xu et al., 2017b). The key genetic factors that impart heat tolerance characteristics along with the genetic factors that impart cold tolerance to rate constants and steady-state Φ_PSII_ observed in P1 and P2, P3 and P4 respectively, could prove to be a quintessential target for breeding climate-resilient crops. These targeted approaches can also be beneficial for tomato growers who can grow tomatoes at lower temperatures which cuts the cost of heating. Brüggemann et al. (1996) have already demonstrated that cold-tolerant tomato lines can be obtained by crossing the cultivated tomato with a cold-tolerant species, *Solanum peruvanium*. Therefore, it is advised to generate a population utilising the genotypes from this study and then identify QTLs (quantitative trait loci) in order to determine which genetic factors influence these traits.

## Conclusion

The study explores the effects of short-term fluctuating light and cold on photosynthesis in four tomato genotypes. Significant genotype-to-genotype variation in photosynthetic responses to both short-term fluctuations and temperature changes was observed; while previous studies have focused on constraints during photosynthetic induction from darkness, this study focuses on the combined effects of cold and fluctuating light on photosynthesis.

The study found that plants’ response to stress, particularly in cold environments, is measured by a decrease in Fv/Fm. Genotype P1 was highly sensitive to cold, however, also recovered at 21°C, and P4 showed the least change. The study found that ΦPSII, a photosynthesis indicator, showed similar trends to Fv/Fm, suggesting it could be a useful indicator of cold effects on photosynthesis without long dark adaptation. Dusk ΦPSII was slightly lower than dawn, possibly due to light treatment or diurnal processes. The genotype P1 had consistently low ΦPSII in all temperature conditions, dropping most during the cold phase and recovering most during the recovery phase.

The study found that factors affecting photosynthesis’s steady-state properties differ from those regulating its response to changing irradiance. Genotypes that respond quickly or slowly to irradiance changes do not necessarily have the highest or lowest ΦPSII. The genotype that responds slowest during step increase does not respond slowest to step decrease. This distinction could help us understand which processes limit photosynthesis under fluctuating light and the genetic factors underlying these constraints. When we compared genotypes, we found that P1’s rate constant for responding to a drop in irradiance remained constant, but that rate of reaction to an increase in irradiance with cold, and even sustained cold exposure, changed less. Similarly, in response to step increases and step decreases in irradiances, genotypes P2, P3, and P4 had ΦPSII that remained largely similar during the cold and recovery conditions.

These genetic factors could be markers for desirable traits to improve photosynthesis under fluctuating light and cold conditions. These traits could be crucial for breeding climate-resilient crops, such as tomato growers who can grow at lower temperatures, cutting heating costs. The study recommends generating a population of genotypes and identifying quantitative trait loci to determine the genetic factors influencing these traits.

## Supplementary figures

**Figure S1.**
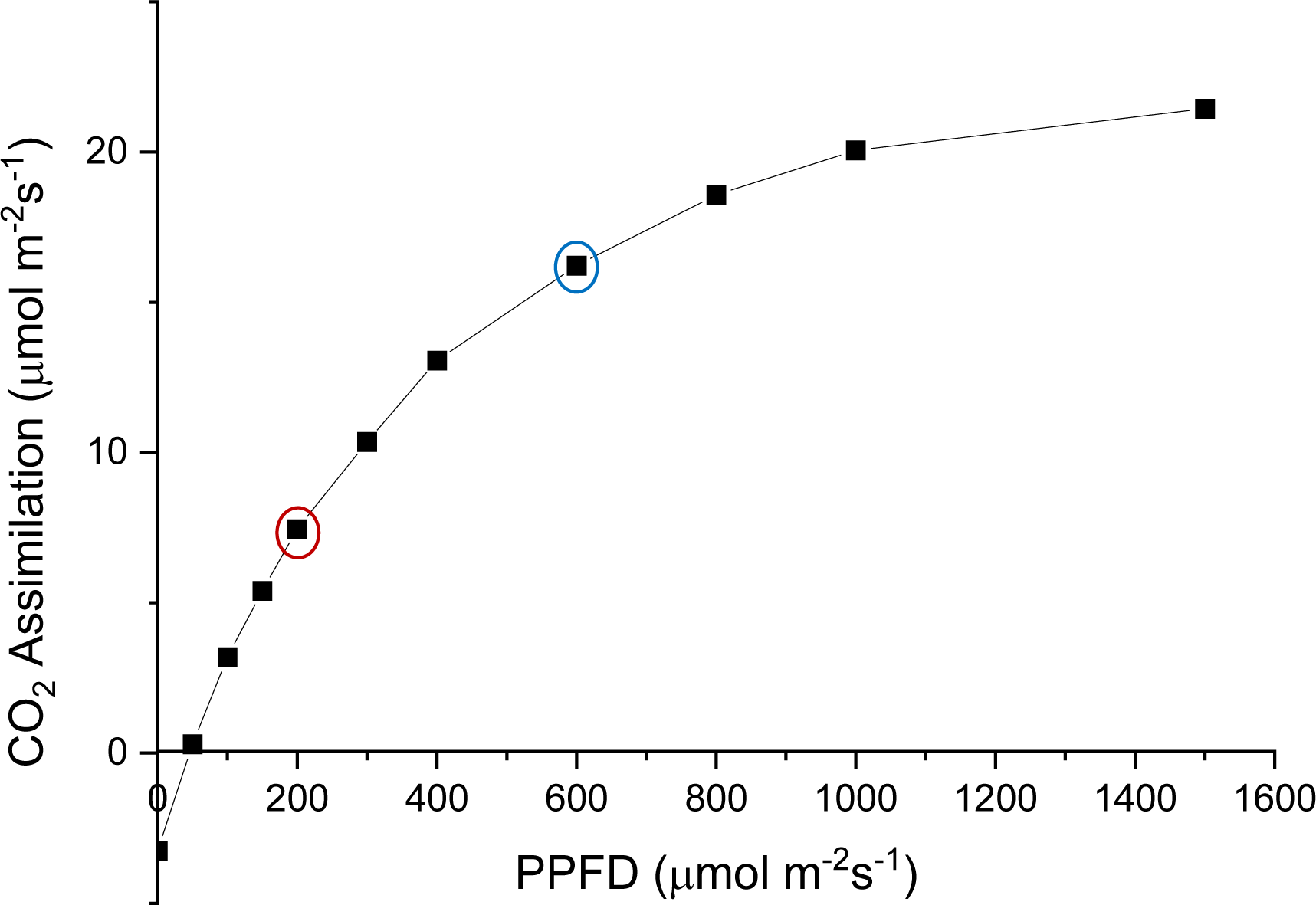
Light response curve for Money Maker grown at 400 μmol m^−2^s^−1^.Reference genotype Money Maker was used to design the irradiance values for step decrease and step increase. 200 μmol m^−2^s^−1^ (red) was taken as the lower irradiance and 600 μmol m^−2^s^−1^ was taken as the higher irradiance (blue).

**Figure S2-.**
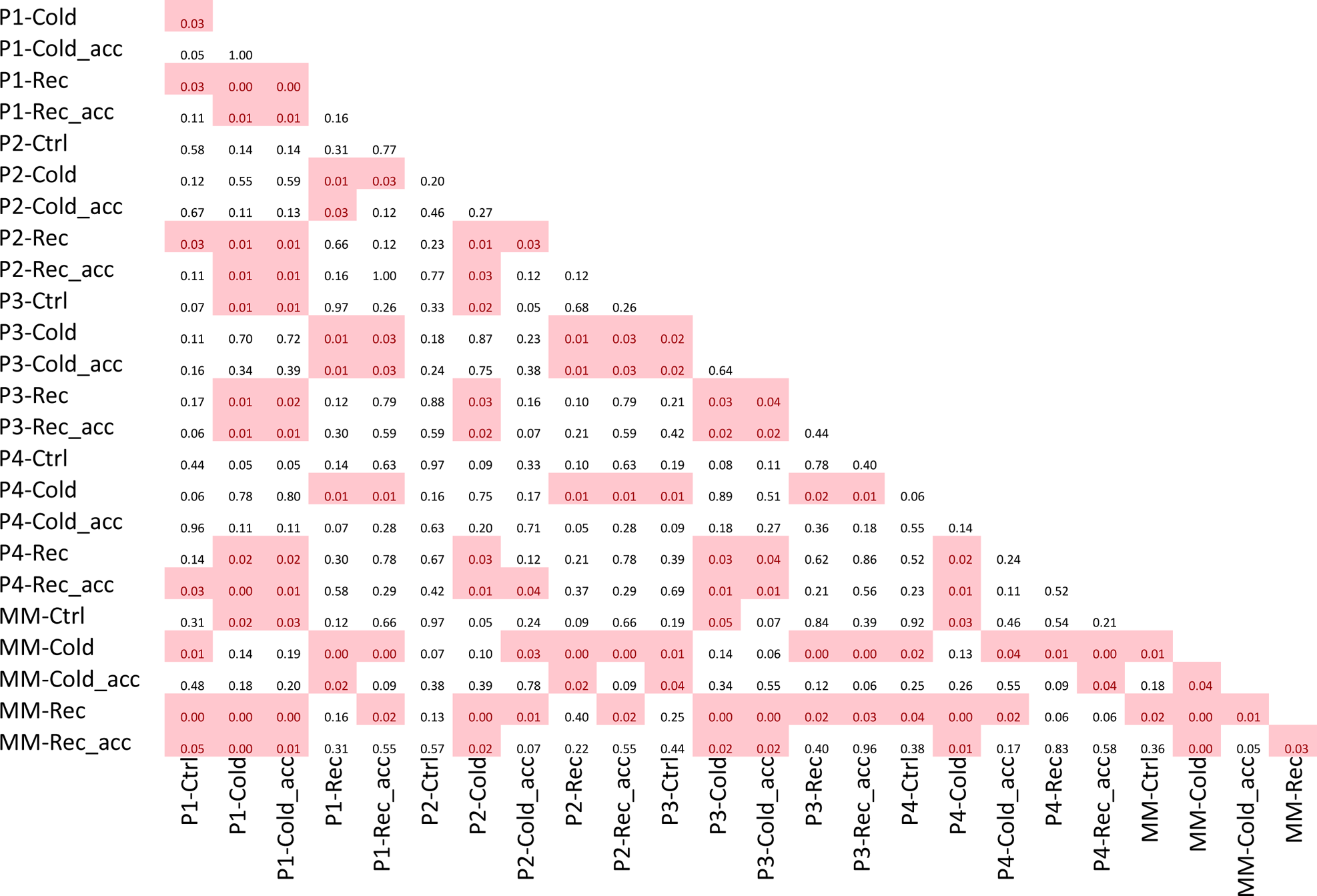
Step increase rate constant significance p-values Welch t-test

**Figure S3-.**
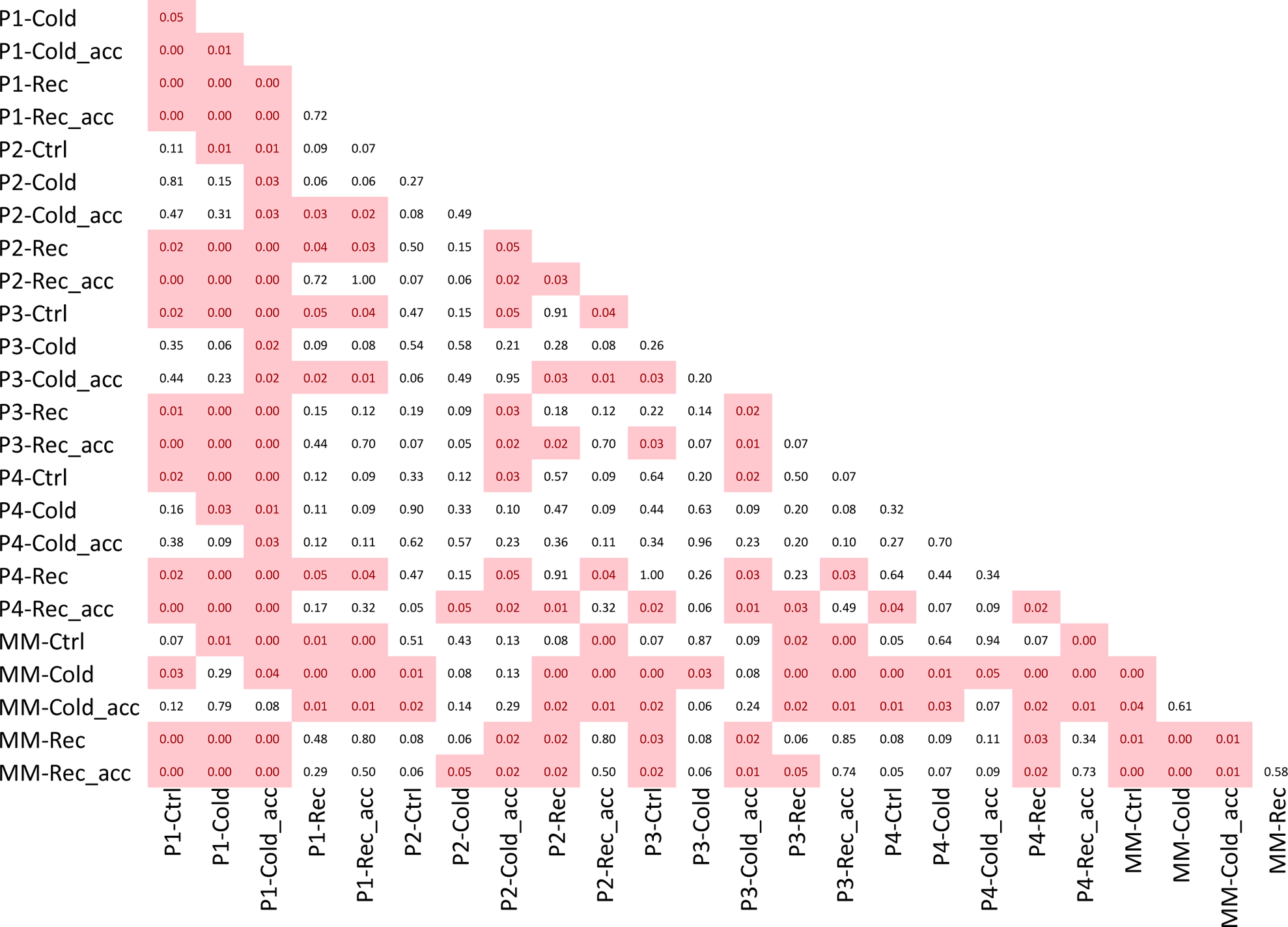
Step increase steady state significance p-values Welch t-test

**Figure S4-.**
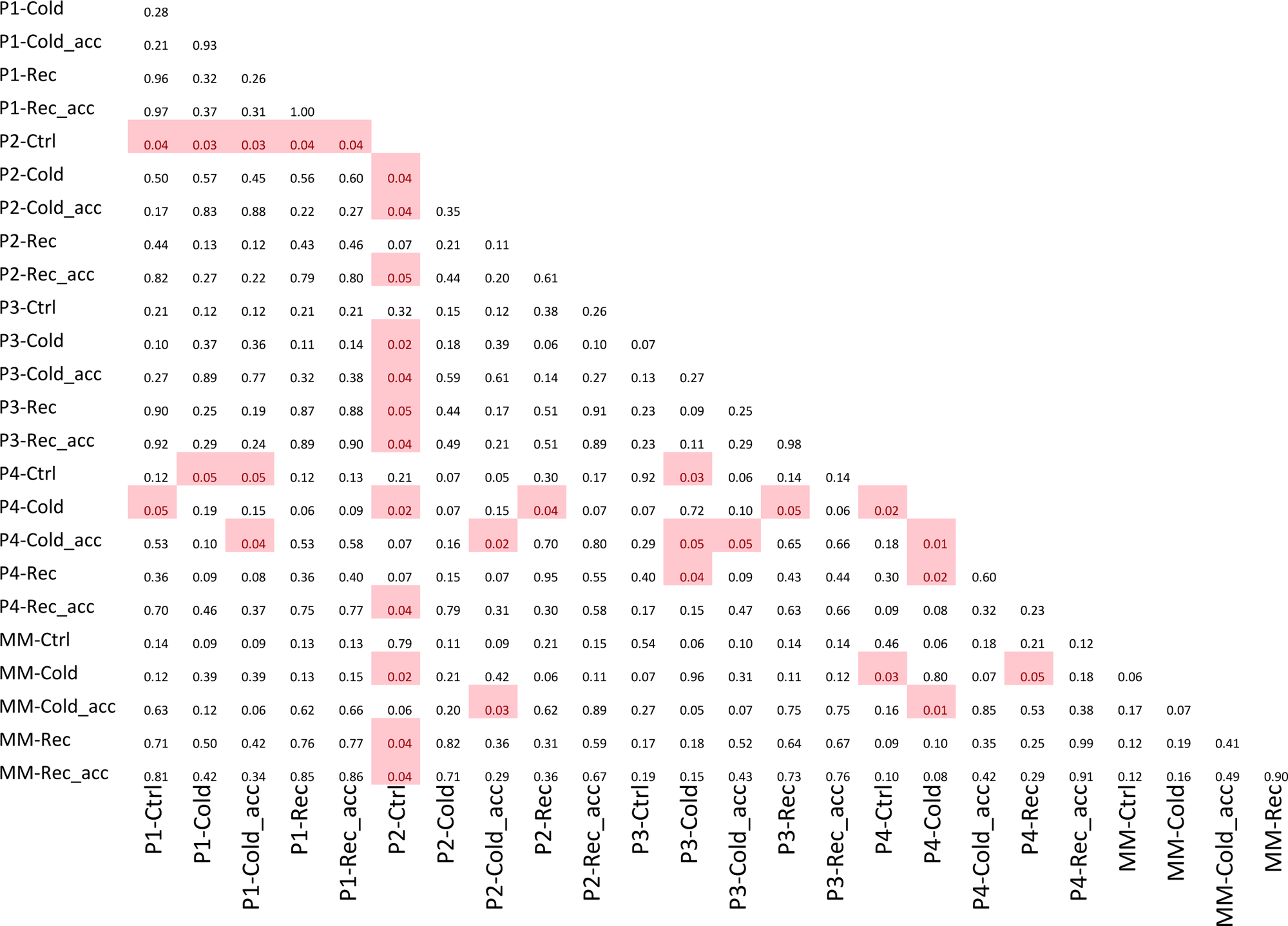
Step decrease rate constant significance p-values Welch t-test

**Figure S5-.**
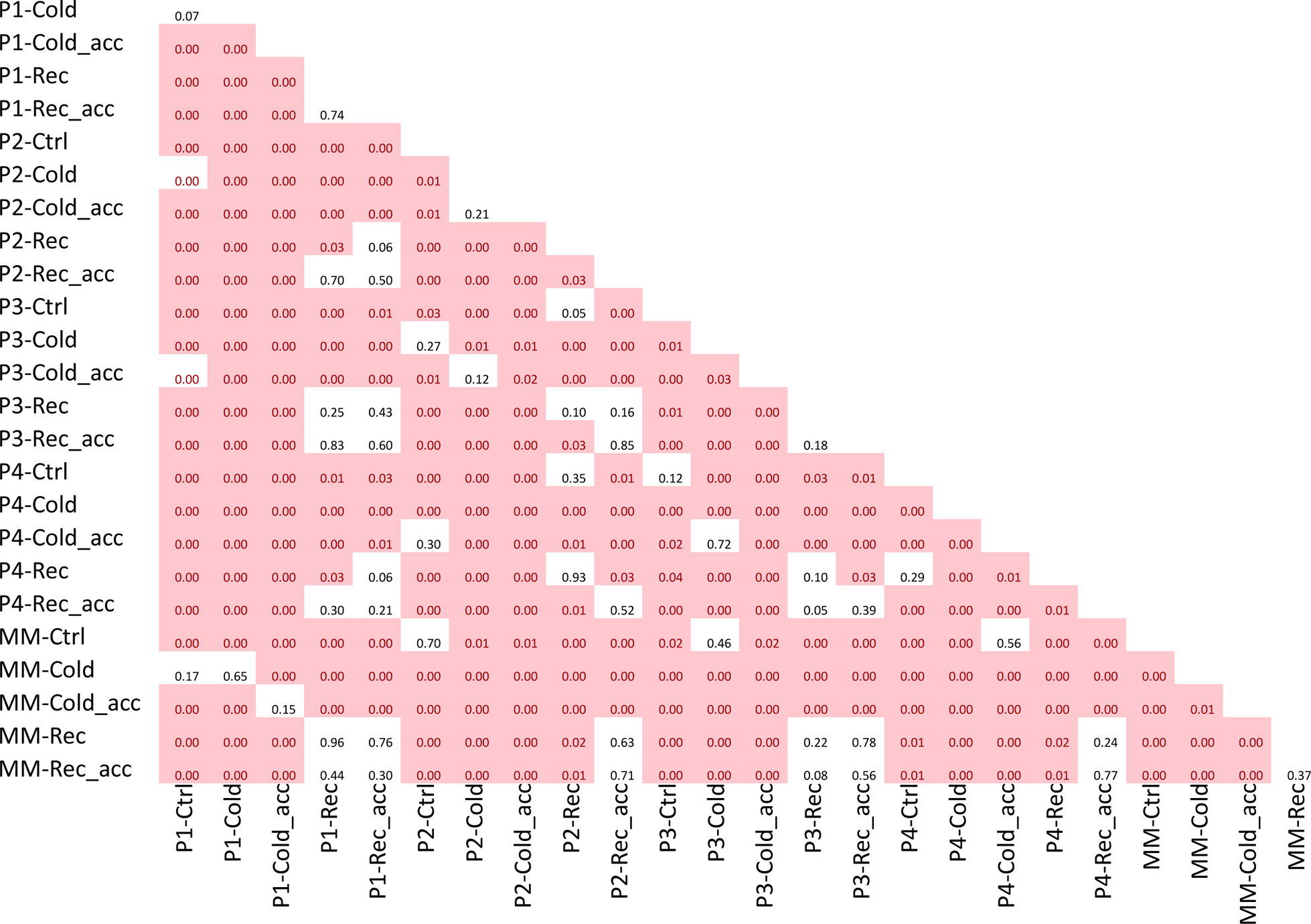
Step decrease Steady state significance p-values Welch t-test

## Acknowledgements

This work was supported by the European Union’s Horizon2020 research and innovation programme (No.862201) project CAPITALISE. We acknowledge Henk Verbakel and Frank Millenaar (BASF Nunhems) for providing tomato seeds. We thank Louise Logie for helping with the phenotyping. We acknowledge the Netherlands Plant Eco phenotyping Centre (www.npec.nl) and particularly Rick van de Zedde Lucas Schmitz, Jannick Verstegen and Jonatan Hovenkamp for assistance in plant phenotyping.

## Author contributions

L.R. and K.J. designed the experiments, performed the research and wrote the manuscript. J.H. and M.G.M.A. supervised the study and revised the manuscript. All authors have read and approved of its content.

## Abbreviations

CO_2_ carbon dioxide

DAS days after sowing

Fv/Fm Maximum photosystem II efficiency

LUE Light use efficiency

MM Moneymaker

Φ_PSII_ Quantum efficiency of PSII photochemistry

